# Hierarchical control of visually-guided movements in a 3D-printed robot arm

**DOI:** 10.1101/2021.08.10.455600

**Authors:** Adam Matić, Pavle Valerjev, Alex Gomez-Marin

**Affiliations:** Behavior of Organisms Laboratory, Instituto de Neurociencias CSIC-UMH, Alicante, Spain; Department of Psychology, University of Zadar, Zadar, Croatia

**Keywords:** robot arm, perceptual control theory, reaching, tracking, human movement

## Abstract

The control architecture guiding simple movements such as reaching toward a visual target remains an open problem. The nervous system needs to integrate different sensory modalities and coordinate multiple degrees of freedom in the human arm to achieve that goal. The challenge increases due to noise and transport delays in neural signals, nonlinear and fatigable muscles as actuators, and unpredictable environmental disturbances. Here we examined the capabilities of a previously proposed hierarchical feedback control model (Powers 1999, 2008), so far only tested *in silico*. We built a robot arm system with four degrees of freedom, including a visual system for locating the planar position of the hand, joint angle proprioception, and pressure sensing in one point of contact. We subjected the robot to various human-inspired reaching and tracking tasks and found features of biological movement, such as isochrony and bell-shaped velocity profiles in straight-line movements, and the speed-curvature power law in curved movements. These behavioral properties emerge without trajectory planning or explicit optimization algorithms. We then applied static structural perturbations to the robot: we blocked the wrist joint, tilted the writing surface, extended the hand with a tool, and rotated the visual system. For all of them, we found that the arm *in machina* adapts its behavior without being reprogrammed. In sum, while limited in speed and precision (by the nature of the do-it-yourself inexpensive components we used to build the robot from scratch), when faced with the noise, delays, nonlinearities, and unpredictable disturbances of the real world, the embodied control architecture shown here balances biological realism with design simplicity.

## 1. INTRODUCTION

Pointing and reaching toward visual targets are nearly effortless human behaviors. However, an explanation of these processes at levels of detail and abstraction that would allow us to build equally capable artificial systems or to treat common disorders in hand and arm control remains elusive. An understanding of such simple motor behaviors should follow from a broader theory of sensorimotor control, while being consistent with the anatomical structure of the underlying system. Such an understanding would provide insights into the origin of laws, invariances, and principles in the behavior of organisms.

The anatomical structure of the nervous system, together with the behavioral analysis of organisms under different conditions and upon different perturbations, suggests that biological control is hierarchical. One of the earliest hypotheses on the hierarchical nature of the nervous system was proposed by John Hughlins Jackson (1884, 1958), discussing the possible evolutionary development of the nervous system in layers (see also Prescott et al, 1999). In turn, neurophysiologist Nikolai Bernstein proposed a hierarchical organization of neural structures underlying movement, where each layer performs a specific function, increasing in abstraction as one ascends the hierarchy (see Profeta and Turvey, 2017). Arguments for the existence of hierarchy of control can also be made from the comparative evolutionary history of the nervous system (Cisek, 2019) and from early development in primates (Plooij and van de Rijt-Plooij, 1990).

However, some findings in spinalized preparations blur the line between the capabilities of different levels in motor control. Cats with a transected spinal cord or cats in a decerebrate preparation can learn to walk on a treadmill (Whelan, 1996); decerebrate ferrets can learn new locomotion trajectories (Lou and Bloedel, 1988, 1992); rats with lesions in the motor cortex can still move in stable, predictable, non-perturbing environments, but not if the environment is rapidly changing (Lopez, 2016). Those experiments show the existence of independent “lower levels” in the spinal cord, capable of relatively complex behaviors on its own, despite normally operating in accord with the higher levels. Therefore, while the consensus seems to be that biological control is hierarchical, it is still unclear what the function of each particular level is, their limits and relationship, or even why there is a hierarchy at all.

Hierarchical architectures have been used in robotics, famously by Brooks in the subsumption architecture (Brooks, 1986). More recently, Merel et al (2019) listed core advantages of hierarchical control appearing in both biological and engineered systems. Hierarchies allow for modularization and simplification of individual controllers and training procedures. Each subsystem can deal with only a part of the incoming sensory information, and, having partial autonomy, can be trained separately with cost functions and performance requirements distinct from the task objective. In contrast, a “flat” non-hierarchical controller would receive and process all the sensory information, directly calculating the behavioral output. Such an arrangement would make the control algorithm both complex and complicated, require extensive training, and result in rather incomprehensible information flows. Thus, at least from an engineering perspective, hierarchical architectures can be very beneficial for adaptive behavioral control.

A prominent normative approach to behavior is optimal feedback control (Todorov and Jordan, 2002a; Scott, 2004, 2012; Shadmehr and Krakauer 2008), which predicts many of the features of human movement and corrects a long-standing bias against the importance of sensory feedback in online movement (e.g. Flash and Hogan, 1985; Uno et al. 1989). However, this theoretical framework does not necessarily suggest a neural substrate for implementation of the proposed control algorithms. In fact, it is still debated whether some features of the optimal feedback control architecture, such as internal forward and inverse models, are computationally necessary, and whether they can be found in the brain or not (Loeb, 2012; McNamee and Wolpert, 2019; Hadjiosif et al 2021). Briefly, forward models estimate the current state from a copy of the motor command and the delayed sensory signals, while the inverse models (also called controllers) provide a motor command that will achieve the desired state given the current state and an inverted model of the plant (Wolpert and Kawato, 1998). The theory does not explicitly address the hierarchical structure of motor control or the role of sensory feedback in subcortical levels and the spinal cord.

Exploring the computational principles that underlie eye-hand coordination and synergistic control in pointing and reaching, William T. Powers designed a series of distinct models of arm control. The first one (Powers, 1999) contained a model of muscles, a binocular vision control layer, and an arm controlled via three degrees of freedom (DOF). The organization of this control system follows roughly the anatomical hierarchical organization of the spinal and some supra-spinal neural structures involved in human motor control and arm coordination. The second model (Powers, 2008) consisted of a 14 DOF arm with more fidelity in arm segment lengths and joint movement limits, but it lacks the muscle model. These models were built by cascading multiple layers of simple proportional and proportional-derivative feedback loops with low-pass filtering. Using hierarchically arranged controllers provides the benefit of avoiding the unfeasible complications of calculations of inverse models, estimates of load properties, and even inverse kinematics.

However, the behavioral capabilities of such simple but powerful models have not been assessed beyond the ideal world of numerical simulations. Our aim here is to test the *in silico* idea *in machina*, namely, to run those arm simulations in a robot arm; to cloth virtual software with actual hardware, thus assessing not only the feasibility of the proposal, but its biological realism in the context of human movement.

Why is this necessary and important? As with many simulations, Powers’ arm models contain idealized representations of the nervous system, the body, and the environment. To name a few: there is no friction, no noise, no long transport delays (although there are some delays), no contact forces, etc. These idealizations are acceptable for initial testing and demonstration of principles, but as argued by Barbara Webb (2001), models of biological structures should be tested in terms of real problems faced by real organisms in the real world. Additionally, as claimed by Brooks (1992), there is a near certainty that programs which work well on simulated robots completely fail in real robots because of the differences between simulated and real-world sensing and actuation. Moreover, designing and building robots that work decently can generate insights about the function of structures in the nervous system that produce analogous behaviors in living organisms (Floreano et al, 2014; Marimoto and Kawato, 2015).

In sum, following this approach, in the present work we adapted and implemented the proposed hierarchical control architecture (Powers, 1999; 2008) to a 4 DOF robot arm in order to examine its theoretical capabilities in the real world (dealing with noise, delays, nonlinearities, and unpredictable environmental disturbances), as well as to generate insights about human control in basic task such as reaching or tracking. In the first part of the manuscript, we show that several fundamental invariant properties found in human hand trajectories — isochrony, bell-shaped velocity profiles and the speed-curvature power law—are also found in the robot arm trajectories without planning or optimization. In the second part, we demonstrate the motor equivalence phenomenon, whereby the robot arm can still perform reaching and tracking movements with a blocked wrist, without learning or being reprogrammed. We also show spontaneous behavioral adaptation to a tilt of the writing surface, to a rotation of the visual field with respect to the arm segments, and to the extension of the robot “hand” with a tool. We conclude by discussing the limitations of both the robot and its control architecture, specifically in the light of modeling fidelity and potentially higher/lower levels of control.

## 2. METHODS

### 2.1. Hardware: The robot arm

We designed and 3D-printed robot arm segments and its rotating base in PLA plastics. Several pictures of the arm and its diagram are shown in **Figure 1**. The robot has four degrees of freedom: shoulder rotation angle, shoulder pitch, elbow pitch, and wrist pitch. They are all actuated via motors M0–M3. The location of the shoulder pitch joint is 4cm above the base level. The upper and lower arms are both 12cm in length, while the hand spans 10cm from the wrist joint to the hand tip. Each joint has a geared DC motor as the actuator and a potentiometer as a sensor to estimate the joint angle and angular velocity. The motors and gear trains come from servos (two HobbyKing 15298 in base and shoulder joints, Futaba S3003 in the elbow, and an N20 DC motor in the wrist). All the electronics and control circuits were removed from the servo motors and replaced by custom control software on the microcontroller. The potentiometers on the gear output shaft were kept to measure the angular position of the joints. The microcontroller used was Teensy 3.1, with a Cortex-M4 processor working at 96Mhz (3.3V logic). It was programmed in C++ in Arduino IDE 1.8.6. It outputs four pulse-width modulated (PWM) signals to two TB6612FNG dual H-bridge 1A motor drivers, each connected to two motors and a 9V 1A power supply, limited to 5V in software. The sampling rate for angle potentiometers and control signal calculation rate on the microcontroller was 200Hz.

**Figure 1.**
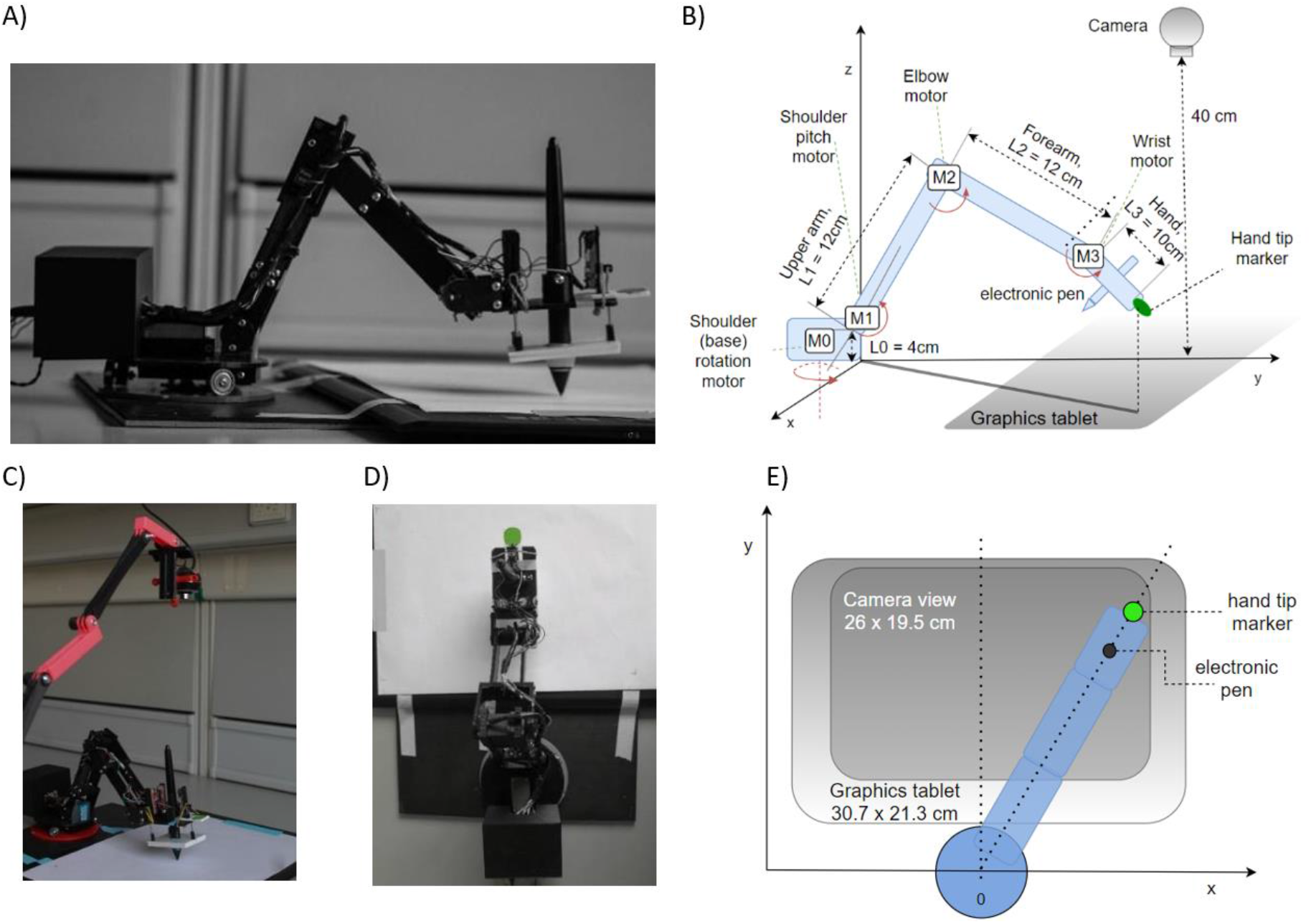
The robot arm system design and implementation. (**A**) Side view showing the body of the robot, enclosed microcontroller, electronic pen and tablet. (**B**) Diagram of the robot arm in perspective view with arm segments L0 – L3, motors M0 - M3, camera, tablet, pen and marker of tip position. (**C**) Photo of the experimental setup, including the top camera. (**D**) Top view photo (camera’s viewpoint), the green circle is used by the visual system as the marker of hand tip position. (**E**) Diagram of the robot from the top view.

To detect and measure the marker position, we used a generic webcam with a resolution of 640 × 480px and a maximum frame rate of 30Hz. The camera was placed at a height of ~40 cm from the writing surface using a 3D printed stand, pointed down toward the marker on the tip of the hand, covering the area of 26 × 19.5 cm, slightly smaller than the graphics tablet active area size. The image from the camera was used to construct two controlled variables, the x and y position of the marker, and formed the basis of visual controlled loops.

To measure pen angle and pressure, we used a graphics tablet Wacom Intuos Pro Paper PTH-860, with an active surface of 30.7cm × 21.3 cm, at a spatial resolution of 0.08mm. The sampling rate of the tablet was 120 samples per second. However, we used 30 samples per second for pen pressure and pen angle control in order to be synchronized with the visual control loops that were limited by the temporal resolution of the camera to 30Hz. The position of the pen as measured by the tablet itself was not used in arm control.

The PC we used for recording and visual processing had an Intel i5 processor, 8GB of RAM, and runs on Windows 10 OS.

We initially designed and placed pressure sensors on the hand of the robot, using three linear sliding potentiometers measuring the stretch of an elastic rubber band when the hand is pressing on a surface (visible on the hand in **Figure 1**). One sensor was placed at the tip of the hand, and two on the base. The sum of travel of all three potentiometers was therefore directly related to the pressure of the palm on a surface, and the difference between the front potentiometer and two back potentiometers was related to the pressure difference and tilt of the hand. The Wacom graphics tablet also reports the pressure of the pen on the tablet and the angle of pen tilt, and these readings proved to be more reliable than our custom sensors and were used in control loops.

### 2.2. Software: The control architecture

Controlled variables in the lower level of the robot arm (**Figure 2A**, blue level) are functions of proprioceptively sensed joint angles and stored arms segments lengths calculated by forward kinematics: the x coordinate of the hand tip in proprioceptive space (xp); the height of the hand tip or z coordinate in proprioceptive space; reach (R), the distance from the base to the hand tip; and δ (delta), the angle of the hand to the x-y plane. These variables were the input to a proportional-derivative (PD) controller with a low-pass filter in controller output, following:

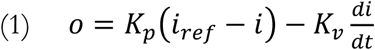

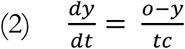

where (1) is the controller equation with controlled (measured) variable *i*, reference input *i_ref_*, output *o*, the derivative of the controlled variable *di/dt*, proportional gain *Kp*, and derivative gain Kv. Equation (2) models the low-pass filter with input *o* and output *y*, and *tc* the open-loop time constant of the low pass filter.

**Figure 2.**
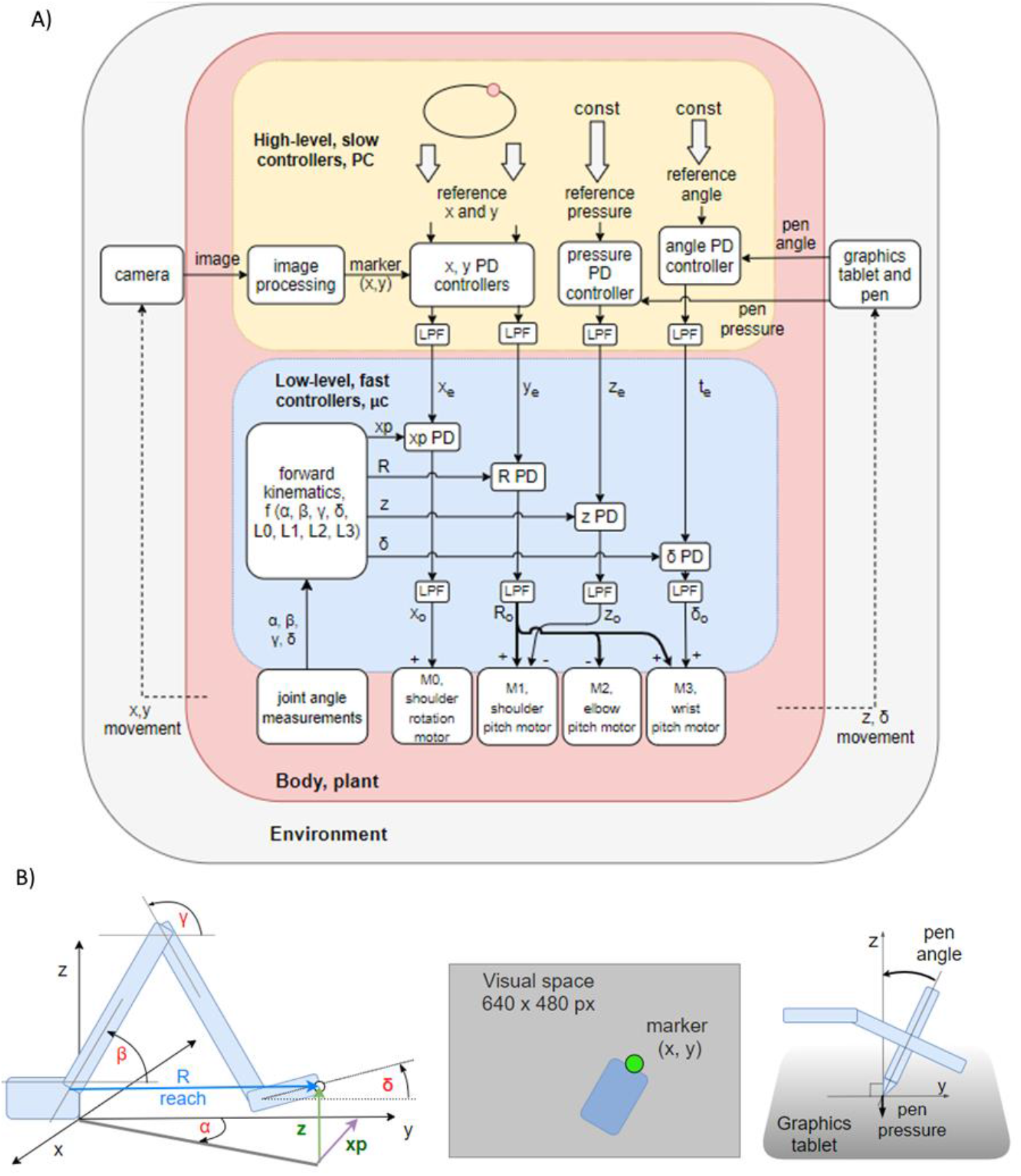
Control diagrams of the robot arm system. (**A**) Block diagram of two levels of feedback loops, high level (yellow) and low level (blue). There are four high-level controlled variables: position of the marker in *x* and *y* dimensions in the visual field, *angle of the pen* to the tablet, and *pressure of the pen* to the tablet. The references are supplied by the experimenter. The outputs from top-level loops are references for the lower-level loops controlling proprioceptive variables: *xp*, the x coordinate of the hand tip in proprioceptive space; reach, the distance from the shoulder base to the hand tip; *z* as the height of the hand tip; and *delta (δ)* as the angle between the x-y plane and the hand. All controllers are proportional-derivative (PD) with a low-pass filter (LPF) in controller output. (**B**) Diagrams showing the geometric definitions of variables in the block diagram, the visual space and a diagram of the pen angle and pressure variables.

The information available to the systems, other than measured variables, included stored arm segment lengths and equations of forward kinematics. Forward kinematics in the input functions were the computationally most complicated part of the lower levels. The outputs x_o_, R_o_, z_o_ and & *δ*_o_ of the lower level low-pass filters are summed to PWM commands to motor drivers (see **Figure 2A**, diagram of lower-level systems). Position in x dimension of the hand tip in kinesthetic space (xp) is controlled by activating the shoulder rotation motor M0. This configuration limits the work area of the arm to less than 180° in the upper half-plane (y>0). Reach R is the distance between the base of the robot and the tip of the hand. It is controlled by simultaneously activating shoulder (M1), elbow (M2) and wrist (M3) motors, with the elbow being activated in the opposite angular direction from the other two. Height z of the hand tip is controlled by moving the shoulder motor M1. Angle delta between the hand and the x-y plane is controlled by moving the wrist motor M3.

All the variables are controlled simultaneously.

For instance, if correcting the height variable creates an error in *δ* angle, it will be treated as a disturbance to the *δ* angle control system and will be corrected simultaneously to the height error. Joint angles are not calculated before starting the movement as in traditional inverse kinematics. Joint motors move until all the errors are reduced. These calculations are performed on the microcontroller (Teensy 3.1, 96Mhz) at 200Hz.

Variables controlled at the top level are x and y positions of the marker in visual space, pen pressure and pen angle (**Figure 2B**). These control systems are implemented on the PC in Python. The image processing algorithm uses the OpenCV library to find the location of a green marker placed on the tip of the hand of the robot (**Figure 2B**). Signal transport delays in visual loops are approximately 180-190ms. The location of the marker is reported in pixels, and it is compared to the reference determined by the experimenter. Each variable is sampled or calculated at approximately 30Hz, determined by the sampling rate of the camera.

We first tuned the lower-level, proprioceptive loops. The tuning procedure started with setting proportional and derivative gains to zero, and the time-constant of the low pass filter to a low value. Next, we increased the proportional gain until oscillations appeared after step references, and then increasing the derivative gain until the oscillations would stop. If the precision were not high enough, then we would increase the time constant and retune the proportional and derivative gains to a higher value, trading bandwidth for precision and stability. The time constant *tc* of the low-pass filters was 80ms for all the controllers at the lower level. For tuning the higher levels, we applied step references and aimed for a critically damped response using the same trial and error procedure as described. The PD controllers at the higher level are identical to lower-level controllers expressed in equations (1) and (2). They differ only in parameters. Loop gains and open-loop time constants are much larger in higher-level loops in order to achieve stability and precision in conditions of large loop delays and noise from the marker location finding algorithm.

### 2.3. Data analysis

Robot hand trajectories were extracted from the camera-recorded positions and estimated hand marker locations. We did not use the position of the pen on the tablet since the position of the hand-tip marker was not identical to the position of the pen. The experimental signals were smoothed with low pass second-order Butterworth filter, with the cutoff frequency specified for each analysis, in order to tame the relatively high levels of noise, aiming for the preservation of position and velocity profiles, and taking into account the speed of arm movement. Trajectories of the computational model were not smoothed.

## 3. RESULTS

Beyond computer simulations and blackboard mathematics, we studied an “embodied control architecture” in the real world (aka, our robot arm) to see how it can deal with tasks commonly performed by humans and other primates, while adaptively managing noise, delays, nonlinearities, unpredictable disturbances, and perturbations.

Having built the robot arm hardware from scratch and having implemented the hierarchical control algorithms as described above, our main goal was two-fold: first, to examine the behavior of the system in its own right, and second, to compare the behavioral features of the robot arm to known properties and invariances of human arm movement.

### 3.1. Task I: straight movements in a reaching task

The first test is a reaching paradigm similar to the center out reaching task (**Figure 3A**) often used in primate and human movement research (e.g. Cisek and Kalaska, 2002). We applied the step reference signal simultaneously to x and y visual tracking loops. There was no central stopping point: for one size of the task, there were 10 movements in each direction, done in sequence, for a total of 40 movements, with a 5-second pause on the endpoints. The task was repeated 4 times with different lengths of movement at 4cm, 8cm, 12 cm, and 16cm (**Figure 3B**). We did not randomize the movement directions, since the robot did not have any learning capabilities that might have influenced the reaction time or movement trajectories.

**Figure 3.**
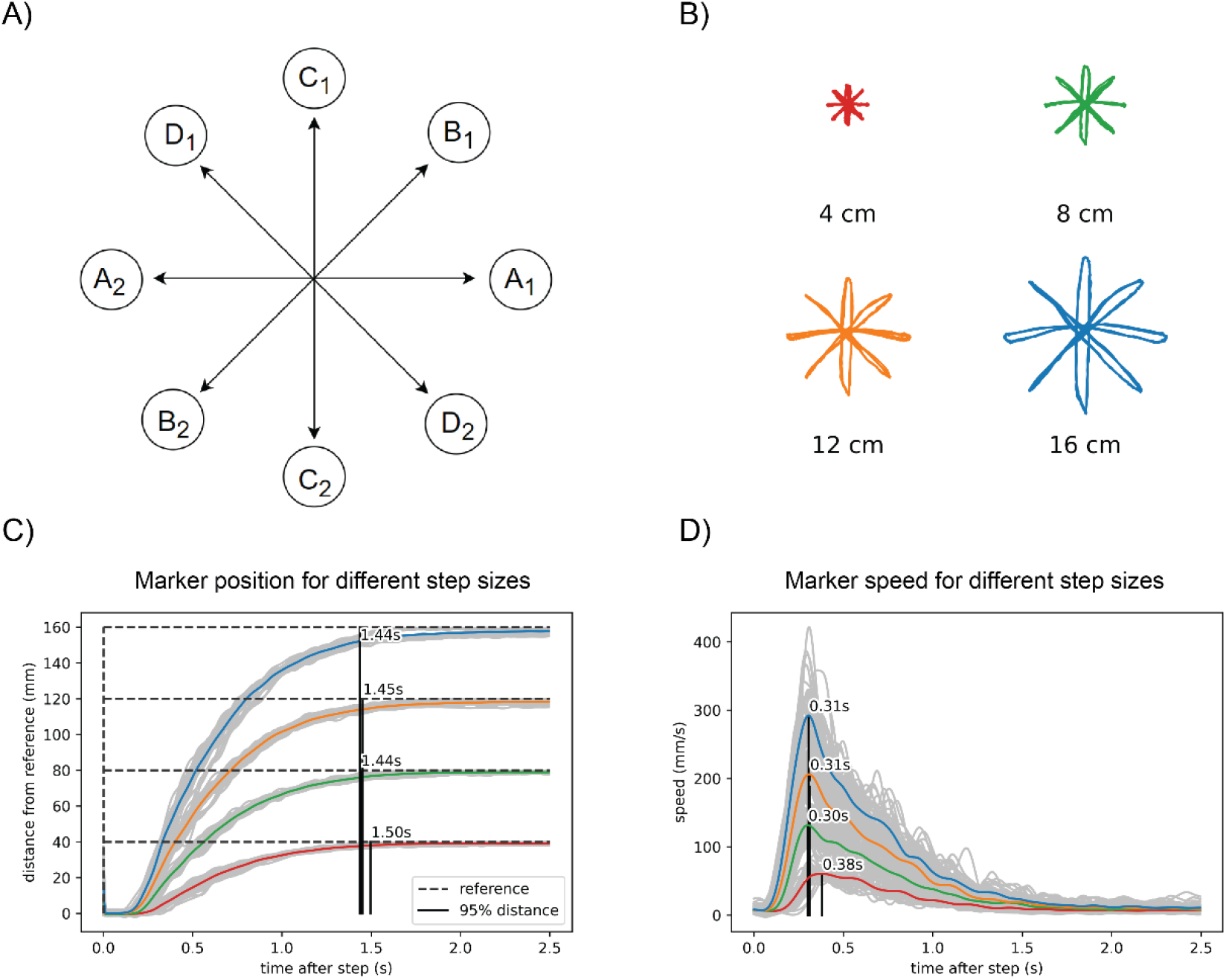
In reaching to a visual reference, the robot arm shows isochrony and bell-shaped speed profiles. (**A**) Task diagram. The reference jumps between points A1 and A2, with a pause of 5 seconds at each point. Then repeats the same pattern at points B1 and B2, C1 and C2, and D1 and D2. (**B**) Marker position data for four different sizes of the reference step. (**C**) Marker position in visual space calculated as the distance from the previous marker position. The marker reaches 95% of the distance to the reference after a step in about 1.45 seconds, regardless of the distance or direction traveled. (**D**) Speed calculated from the same data shows a bell-shaped profile, scaled in height to step distance. The position data in plots B and C and was low-pass filtered with a 2^nd^ order Butterworth filter with a cutoff at 15Hz, while the speed data filter had a cutoff at 5Hz.

We found that the robot performs straight movements across different lengths and different directions in approximately the same time: 1.45 seconds (**Figure 3C**). For the shortest movements (4cm) there was a deviation of 50ms from the average duration of longer length movements. The speed profile (**Figure 3D**) was roughly bell-shaped with a shorter rise and longer fall segment and scaled with the length of movement. In all movements, the maximum speed was achieved at approximately the same time after the reference step (peak at 0.33s, **Figure 3D**), except in the shortest movement of 4cm, where the peak of maximum speed was 70ms later than the average of the other movements. Robot trajectory data were low-pass filtered using a second-order Butterworth filter with a cutoff at 3Hz.

### 3.2. Task II: curved movements in a tracking task

Producing elliptic traces or drawings in humans in a fast and fluid manner results in a speed-curvature relationship known as the 2/3 speed-curvature power law (or 1/3 power law, depending on the variables used), first described by Lacquaniti and colleagues (1983). In the second task, we tested the production of curved movements. We used a continuously moving reference point, and we report on the situation where the reference moved at a constant speed along an elliptic path. An elliptic trajectory with a constant tangential speed is a non-power law trajectory (β≈0, r^2^≈0); the x and y components are not pure sinusoids, but also contain higher frequency components. In the low frequencies, the speed of the robot was close to the reference speed, but at higher frequencies, the speed of the robot was not constant and had a sinusoidal profile and a lower average than the reference speed. At the highest frequency of input (f = 0.826Hz, speed ≈ 406mm/s), the output trajectory followed a speed-curvature power law with an exponent of β ≈ - 1/3 and r^2^ = 0.78. (**Figure 4C**).

**Figure 4.**
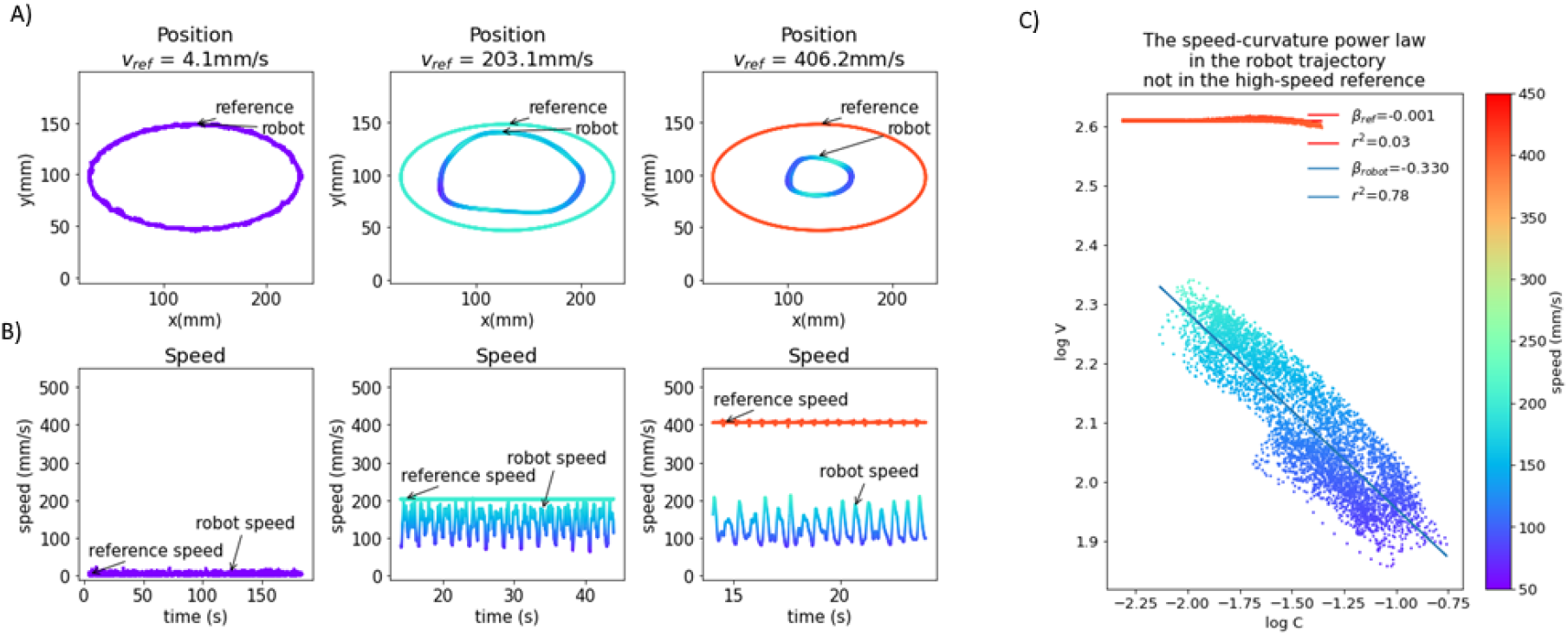
The emergence of the speed-curvature power law at high speed. (**A**) Reference and robot positions. When the reference speed is low, the position of the robot hand tip marker in visual space matches the speed of the reference. At higher speeds, there is a magnitude attenuation – the ellipse shape is smaller. (**B**) Plot of reference and robot speeds. At low speed, the robot speed matches the reference. When the reference speed is high, the speed of the robot is lower and oscillates, even though the input speed is constant. (**C**) Oscillations in robot speed for the highest speed input are regular and correlated with local curvature of movement, following the speed-curvature power law with the exponent of β≈-1/3 and r^2^=0.78. The reference speed is constant (400mm/s), and there is no power law (β≈0, r^2^≈0). In all the plots colors indicate speed, as shown in the color bar on the right. Position data was smoothed with a 2^nd^ order Butterworth filter with a 3Hz cutoff.

This seems to support the hypothesis that the power law is a consequence of the physical limitations of the human arm, and not a planned invariance. However, the size of the drawn shape was smaller than the reference shape because both x and y components of the reference are attenuated in the output. To further probe the question of the origin of reaching and tracking invariances, and to minimize the effect of noise, we created and fitted a model of visual loop behavior.

### 3.3. *In silico* modeling and characterization of the *in machina* system

The system controlling the visual y variable (**Figure 5A**) is nonlinear because increasing reach affects the *y* position differently depending on the angle of rotation of the base (α). The three motors involved in changing reach (Fig. 5A) are different in power and mechanical linkage; they have a different effect on changes in reach depending on the position they are in. Despite nonlinear elements, the behavior of high-level visual loops can be described fairly well by a set of linear second-order equations, modeled in the block diagram (**Figure 5B**), the free-body diagram (**Figure 5C**), and equations:

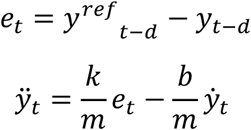

where *t* is time, *e* is position error, y_ref_ the position reference, y position, d is loop transport delay. The values of the coefficients used are k/m=40, b/m=27.5, and d=0.185s, which were found by fitting the behavior of the model to the behavior of the robot in the step reference task with a 12cm step distance. The best fit values indicate a slightly overdamped second-order system. We modeled the x and y control loops as independent systems with equal parameters.

**Figure 5.**
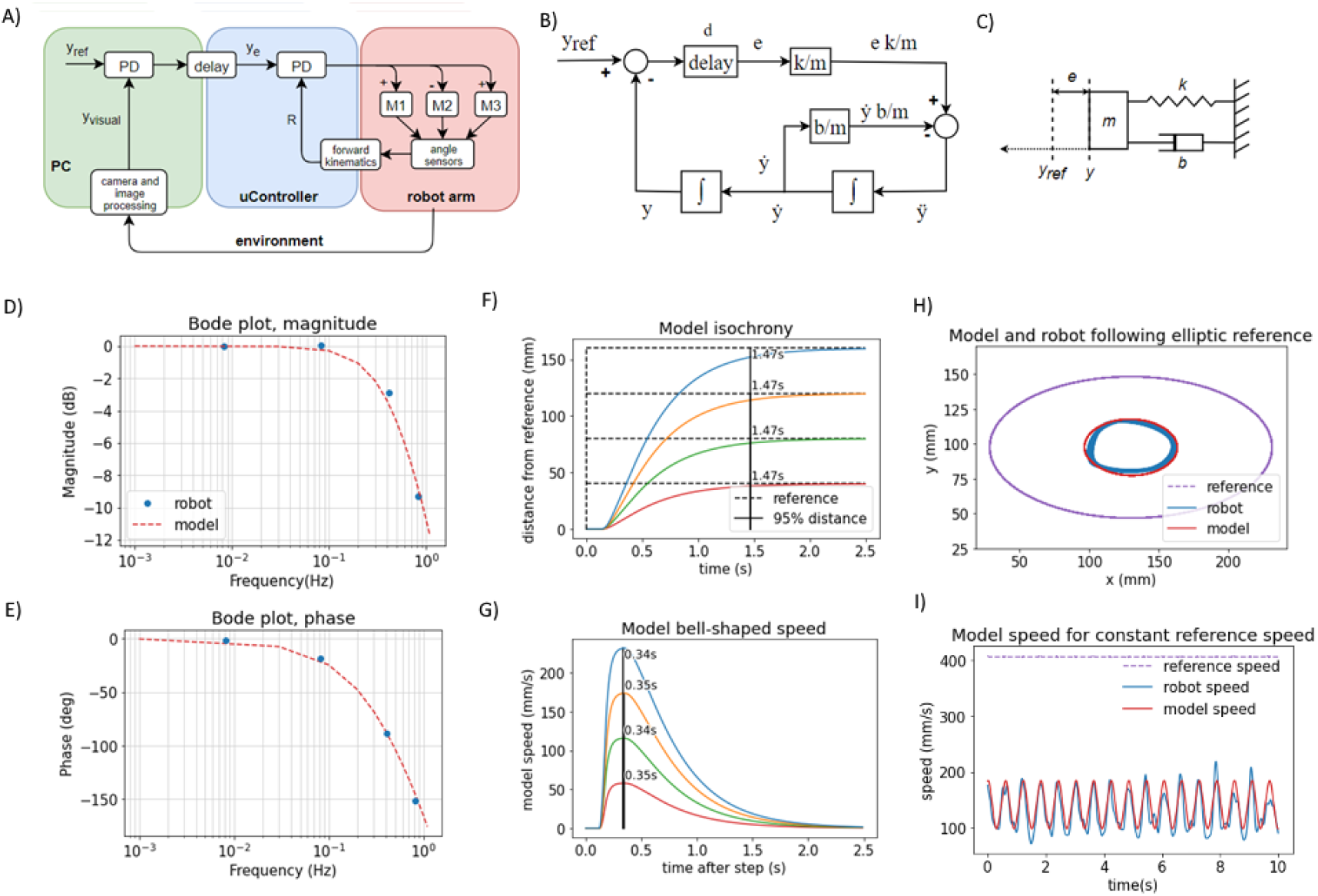
The behavior of visual feedback loops can be represented by a second-order model. (**A**) Diagram of the elements in the control of the position of the hand in visual space in y dimension, including the lower-level control of reach distance. (**B**) Block diagram of the model with variables, parameters, and functions. (**C**) Free body diagram of a mechanical setup analogous to the model mass on a spring with damping. (**D**) Bode magnitude plot shows how the amplitude of output signals at high frequencies is attenuated in relation to the input signals. (**E**) Bode phase plot showing the frequency-dependent phase difference between the input and the output. The model reproduces the frequency-dependent properties of the robot’s visual loop. (**F**) The model produces a similar pattern of isochronous movement independent of distance in the step reference task; as well as (**G**) bell-shaped speed profiles in the same task (**H**) The model produces an ellipse smaller than the reference ellipse, similar to the robot, due to attenuation of high-frequency inputs. The frequency of the input here is 0.826Hz, and the attenuation is approximately 9.5dB. (**I**) In the same task, the speed of the model follows the same sinusoidal pattern as the speed of the robot, even with the constant speed of the reference. Model trajectory was not smoothed in any of the plots.

This model is analogous to a mass-spring-damper system (**Figure 5C**) with a movable equilibrium position and a pure delay element. The approach trajectory of the marker on the hand of the robot to the visual reference in the step-reference task is similar or analogous to the approach trajectory of an object of mass *m* on a spring with stiffness *k* and damping *b* toward its equilibrium position y_ref_, where the displacement of the equilibrium position happens after a delay of *d* seconds. The spring constant is an analog of the visual gain or sensitivity to error; the damping coefficient b is an analog of the combined effect of visual velocity gain (damping term in the visual PD controller), gains at the proprioceptive level, and friction between the pen and the tablet; the mass in the mass-spring-damper system is an analog of the combined contribution of the mass of the robot arm, time constants in the visual loops and inertia of electromotors; and the delay is the total duration of the travel of the signal around the visual loop, combining camera latency, frame rate and transmission delays of the serial protocol between the PC and the microcontroller.

We examined the behavior of the robot in response to sinusoid inputs across a range of frequencies (0.008 Hz, 0.083 Hz, 0.413 Hz, 0.826 Hz). We applied input as the visual reference signals and measured the output as the position of the marker in the visual field over time. We calculated the relation of output amplitude A_out_ to input amplitude as 20 log_10_ (A_out_/A_in_) for the Bode magnitude plot, and we calculated the phase difference between the input and output sinusoids for the phase plot. We then interpolated the plot using the second-order mass-spring-damper model (**Figures 5D & 5E**, model interpolation in red, experimental values in blue).

Looking at the Bode plot (**Figures 5D & 5E**), we can see that the system is stable for all input frequencies. At the gain crossover frequency of approximately 0.1Hz, the system has a large phase margin of about 160°, and at the phase crossover frequency at approximately 1Hz the magnitude attenuation is 11dB, which satisfies the stability criterion of having both the phase and gain margins positive. The bandwidth is limited by the large transport and processing delays in the visual loops, approximated at 185ms. The delays cause a phase shift that takes an increasingly longer part of the sine period with the increase in frequency and are compensated by low pass filtering the controller output (see Equations 1 and 2 in Methods section).

We repeated the step-reference task with the model (**Figures 5F & 5G**). The duration of movement is isochronous across different distances: it takes the same amount of time, 1.47s, to cross 95% of the distance to the reference. The speeds are bell-shaped and scaled with distance, but they all reach a peak after 0.37s, replicating very nearly the behavior of the robot arm.

Finally, in the ellipse tracking task (**Figures 5 H & 5I**) at the highest frequency of 0.826Hz of input, the model replicated the size of the robot trajectory, and also the properties of the speed profile. Even when the reference speed was constant, at this frequency, the speed profile of the model was sinusoidal. Model trajectory followed a speed-curvature power law with the exponent β=0.40, r^2^=0.98.

Model trajectories in panels (**Figure 5D** to **5I**) were produced by a model with the same, constant coefficients, simulated with a time step of 5ms and Euler integration. Model trajectories were not low-pass filtered.

### 3.4. Task III: Adaptation upon blocking the robot arm’s wrist joint

The hallmark of biological motor control is robustness to perturbations. In further testing of the robot arm, we applied different perturbations to the controlled variables, keeping them constant for the duration of the task and not changing any of the parameters in the controllers or other parts of the software of the robot. We blocked the wrist, tilted the writing tablet, added a tool that extended the arm, and rotated the visual field.

In this trial, we blocked the wrist joint and compared the performance of the robot in visual tracking tasks to the performance in normal operation where the wrist was moving. Without any changes to the code or parameters of the control systems, the robot arm performed the tasks even with the wrist blocked. In normal operation, the variable *reach* (**Figure 6A & 6B**) is affected by three motors – in the shoulder (M1), elbow (M2), and wrist (M3). The time-plot of the variables in normal operation in the step reference task (**Figure 6D**) shows joint angles that illustrate how all three motors contribute to the movement. When the wrist is blocked, reach is maintained at the same desired value as in the normal situation. However, reach is not affected by three motors, but only by two: the elbow and shoulder motor automatically pick up or compensate for the work normally done by the wrist motor because their activation is proportional to the reach error.

**Figure 6.**
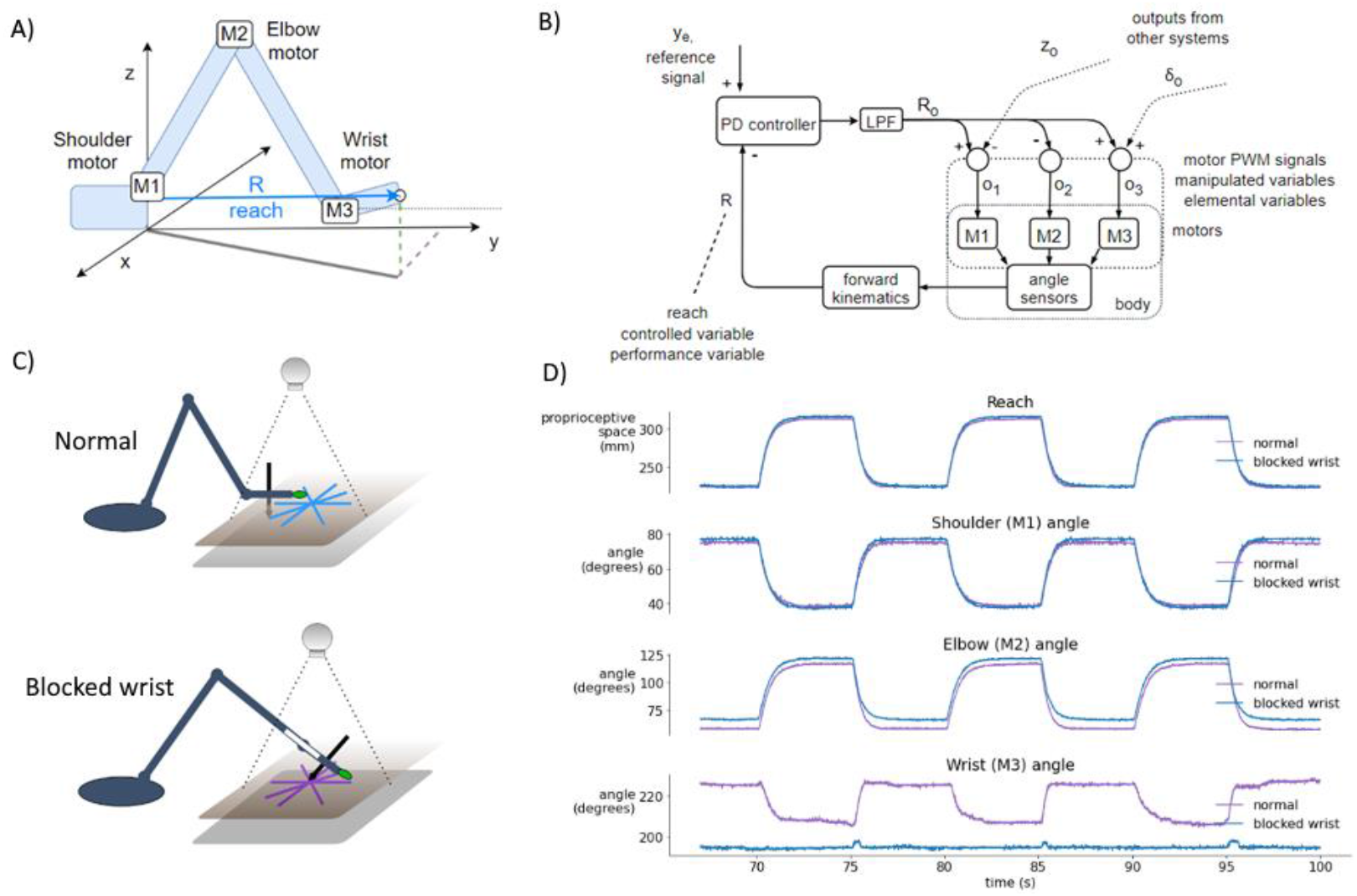
Blocking the wrist is compensated by the reach synergy. (**A**) Geometric definition of *reach* - the distance from shoulder base to the tip of the hand, and locations of joint motors. (**B**) Block diagram of computational and physical processes in the reach synergy, with marked performance and elemental variables. (**C**) Diagram of the normal and blocked wrist setup. In the normal setup, the wrist moves freely, keeping the hand parallel to the tablet (and the pen perpendicular). In the blocked-wrist setup, the motor of the wrist is not powered, and the wrist is locked at ~180° to the forearm. (**D**) Segment of a reaching task with a step reference, comparing the normal and blocked-wrist situation. With the wrist blocked, reach is nearly identical to reach in the normal situation, but shoulder and elbow motors are activated more to compensate. Shoulder and especially elbow angle show differences in both situations.

The block diagram shows the flow of information in the reach synergy (**Figure 6B**). We can describe the system using the terminology of Latash (2008) or Latash and colleagues (2007): the performance variable is *reach*, and it is maintained at its reference level y_e_ by varying elemental variables – activations o1, o2 and o3, of motors M1, M2 and M3, respectively. The activations are calculated by weighting the output R_o_ of the controller, and summing the signals with outputs from other systems, here control of height z and wrist angle δ. These sums (o1, o2 and o3) are used as activations of motors, as pulse-width-modulated signals from the microcontroller to the motor driver chip.

### 3.5. Task IV: Further perturbations: tilting the world, using tools, rotating point-of-view

We applied static disturbances or perturbations to controlled variables either directly or indirectly to examine the adaptiveness and robustness of the robot control architecture. In the normal condition, without additional perturbations, (**Figure 7A**) the writing tablet is horizontal, the wrist is mobile, the marker is on the tip of the hand of the robot, and the visual coordinate system is roughly aligned with the proprioception coordinate system. The angle of the pen to the tablet is sensed and maintained at 0 degrees (pen is perpendicular) by moving the wrist, and the pressure of the pen to the tablet is sensed and maintained at or near 50%.

**Figure 7.**
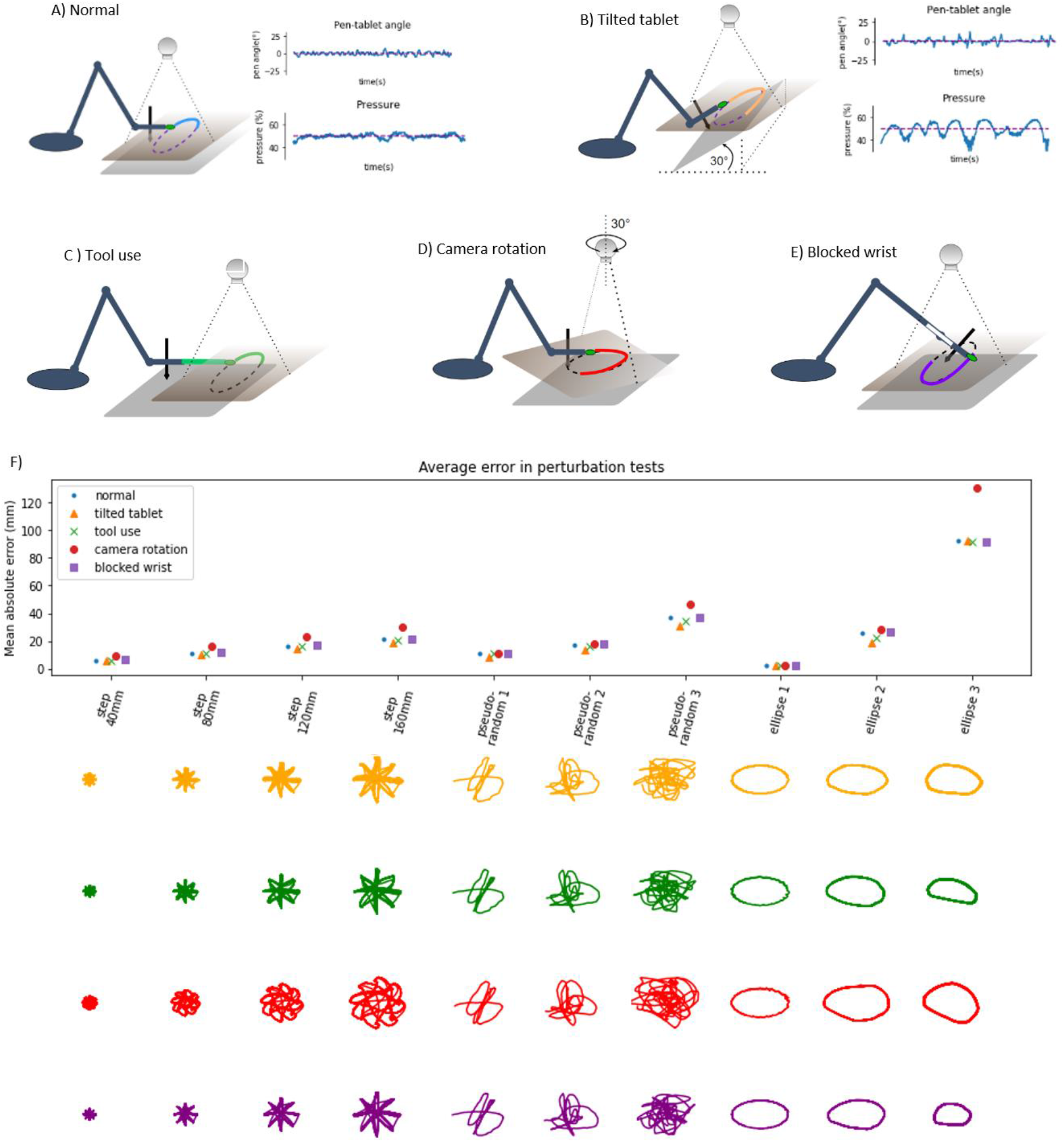
Diagrams of perturbation conditions and a plot of tracking performance in different tasks. (**A**) the normal condition diagram and plots of pen angle and pressure from a pseudorandom tracking task, reference in red and measured variable in blue. (**B**) The tablet is tilted 30°, distal part is lifted, plots show the pen angle and pressure in a pseudorandom tracking task, reference in red and measured variable in blue. (**C**) Diagram of the tool use task, a 12cm plastic piece is attached to the hand of the robot, and the maker placed on the end. (**D**) Diagram of the camera rotation task; the camera is rotated by 30° when compared to the ‘normal’ conditions. (**E**) Diagram of the blocked wrist. (**F**) Average distance (absolute error) between the marker and the reference in the visual space, across different conditions and tracking tasks. Bellow the plot are robot trajectories for each perturbation condition and task.

In the tilted tablet condition (**Figure 7B**), the end of the tablet distal to the robot was lifted to an angle of 30°. This perturbation challenged the pen pressure and pen angle control systems because the pressure control system needs to continuously modify the height of the hand in order to keep the pressure at 50% and still move toward the reference in the visual space. The pen angle control system needs to modify the wrist angle so that the pen is always orthogonal to the tablet surface. The plots of pen angle and pressure show that those variables were maintained near their reference values despite the perturbation, with somewhat more error than in the normal condition.

In the tool use task (**Figure 7C**), we added a 12cm long plastic piece to the tip of the arm and moved the marker forward to the end of the plastic piece, creating a situation resembling tool use, as now the tip of the ‘tool’ was tracking the reference. We moved the camera about 12cm forward to keep the workspace in the visual field. The robot performed the task without learning or reprogramming visual transformations. In the next task, we rotated the visual field by 30° (**Figure 7D**) by rotating the camera and keeping the robot in place. This amount of rotation is near the limits of performance – the robot performed the task with higher amounts of error, visible on the patterns (red) on the plot.

The perturbations summary plot shows the average absolute error as the average distance of the hand marker from the reference in all perturbation tests across different tasks. The error grows with distance or the size of the step in the step-reference task. In pseudorandom tracking, the error grows with ‘difficulty’, where more difficult tasks have a higher magnitude of high-frequency signals. In the ellipse tracking task, the error grows with frequency or with the speed of the reference. Thus, our robot arm is highly robust to external perturbations, akin to human movement.

## 4. DISCUSSION

We have demonstrated how and explained why a custom-made robot arm (**Figure 1**) with a hierarchical control architecture (**Figure 2**) based on simulations by Powers (1999, 2008) displays basic features characteristic of biological movement. Being robust to noise and delays, the robot’s behavior complies with isochrony and displays bell-shaped velocity profiles in a reaching task (**Figure 3**). In tracking a moving target, the robot complies with the so-called speed-curvature power law of human movement at high speeds (**Figure 4**).

We must acknowledge that the proposed control architecture does not fully explain the production of reaching and tracking trajectories since (i) the reaching trajectories of the robot are isochronous for different reach distances, while humans may change the reaching duration according to speed or accuracy demands of the task (Fitts 1954); and (ii) the tracking of elliptic reference trajectories does not recapitulate the geometrical trace or exact speed profile of the reference, with the amplitude of movement falling with frequency, and the shape reducing in size.

However, it is clear that the hierarchical organization of such a control system affords a lot of flexibility to the robot arm even without learning algorithms or online optimization. Moreover, we have demonstrated adaptive behavior to structural perturbations such blocking the robot’s wrist (**Figure 6**) and to environmental configuration disturbances such as tilting the writing tablet, extending the hand with a tool, or rotating the visual field (**Figure 7**). The main reason for such a robust behavioral emergence seems to be the choice of the controlled variables and the hierarchical arrangement of their control systems. We further discuss these findings below.

### 4.1. Frequency response and stability despite noise and delays

The frequency response of the robot’s visually guided behavior (**Figure 3**) shows how the arm behaves in response to input signals of different frequencies, where the “input” is the reference or setpoint input to the hand tip position in the visual space, and the “response” is the hand tip position in visual space. We can see that the system is stable for inputs of any frequency because both the phase and gain margins are positive. The system has a slightly overdamped second-order response (simplified model of the robot arm discussed later in more detail), and it acts as a low pass filter with the gain crossover frequency of approximately 0.08–0.1 Hz. This means that magnitudes of all input signals above this frequency will be attenuated.

Long transport delays are often cited as the main reason for relatively complex delay-compensation schemes such as forward models (e.g. Miall and Wolpert, 1993; Kawato, 1999; Desmurget and Grafton, 2000). However, we show that an alternative, simpler scheme might work. Delays in feedback loops cause a frequency-dependent phase shift so that at high frequencies of input, the phase shift might cause the actuator action to add to the error, creating positive feedback as opposed to negative feedback where the error is reduced, which may cause instability and oscillation. Transport delays in the human visuo-manual loops in tracking pseudorandom targets are approximated at 100–150ms (Viviani et al 1987, Parker et al, 2017). Visual loops of the present robot also contain large signal transport delays, 180-200ms, that come from camera latency and refresh rate, as well as serial protocol communication delays between the microcontroller and the PC. A forward model, given the efference copy of the motor command, estimates the state of the arm at current time, instead of waiting for the delayed feedback signals.

Here we show that stability can be achieved more simply by reducing the bandwidth of the system—trading the bandwidth for stability—using low pass filter elements in the outputs of controllers (**Figure 2**). This maintains the visual loop gain high when the input frequency is low and reduces the effective gain for high-frequency inputs to avoid positive feedback (Powers, 2008). Additionally, the ‘real world’ is highly unpredictable, and various disturbances acting on the arm would invalidate any prediction made by the forward model that takes only the motor command into account. The present architecture avoids the problem by always using feedback signals, affected by both motor action and environmental disturbances, as representations or parameters in calculation of actual state.

The robot arm is still capable of producing movements with high peak speed (**Figure 3**), while movements with both a short duration and a high peak speed might be produced if the movement was stopped by an obstacle. This might be a mechanism involved in fast, short movements of human hands in e.g. pressing piano keys.

Most of the effects of sensory noise in the high-gain visual level systems seem to be averaged over time by the low-pass filter and do not affect the movement, especially at low speed, and don’t require additional compensation mechanisms. In the lower levels, sensory noise does not affect movement because the gains are low.

### 4.2. Isochrony, bell-shaped velocity, and the speed-curvature power law

In humans, isochrony was found in drawing figures of different sizes (Viviani and McCullom, 1983; Viviani and Flash, 1995). It was also found in macaques in natural settings (Sartori et al, 2013), but not consistently in laboratory settings (Castiello and Dadda, 2018). In the Fitts tapping task, the time of movement is related to the so-called index of difficulty, and isochrony is present not for all movements, but for tasks of the same index of difficulty (Guiard, 2009). The tapping task illustrates the speed-accuracy tradeoff: the faster we move, the less accurate our movements will be. Then, in order to preserve accuracy, presumably, when aiming for smaller targets, we slow down our movements.

We have also found that the robot performs isochronously all movements in the step-reference task, regardless of travel distance or direction (**Figure 3**). There were no accuracy requirements, but we found that increasing the frequency and speed in the ellipse tracking task decreased the accuracy, suggesting a speed-accuracy tradeoff in the movement of the robot. This tradeoff seems to be caused by several factors: (i) the low-pass filtering properties of the arm, resulting from its inertia, relatively low power of the actuators and also explicit low-pass filter elements, and (ii) the increased influence of lower-level nonlinearities and control system interactions on the behavior of the robot because the errors on the lower levels were not corrected fast enough.

In humans, during rapid straight-line hand movements, the speed profile is not constant. The movement starts slow, accelerates to a point, then decelerates to a stop, forming a bell shape over time or distance. Some researchers report symmetrical bell shapes where the peak speed is in the middle of the movement (Flash and Hogan, 1985), and some report less symmetrical profiles (Soechting, 1984). In the step-reference task, we found that the speed profiles are bell-shaped, asymmetrical with a short acceleration phase and longer deceleration, and with the maximum speed scaled with the extent of the movement (**Figure 3**). We interpret this speed profile emerging as a consequence of programming visual level loops as proportional-derivative controllers. Since there was no trajectory planning or online optimization, our results show that it is possible to achieve such profiles with a simpler control architecture.

The speed-curvature power law with a two-thirds coefficient is observed in rapid elliptical movement in humans. This phenomenon can be roughly described as movement at lower speed in areas of high curvature and relatively higher speed in areas of low curvature (Viviani and Terzuolo 1982, Lacquaniti et al. 1983). The production of a power law trajectory is not obligatory in principle because the hand might take many of the infinite possible trajectories along the same path. However, the set of possible trajectories is limited by the physical properties of the hand and the environment. For instance, the hand will never move instantaneously from point A to point B, as there is a limit to the force produced by muscles.

Remarkably, when the robot was tracking a high-frequency elliptic reference, we found the speed-curvature power law in the measured movement, even when it was not present in the reference input. This result is consistent with the optimization of jerk in the movement planning phase (Viviani & Flash, 1995; Huh & Sejnowski, 2015). However, in our case we would be getting an optimal jerk trajectory “for free”, again without an explicit optimization algorithm. This result shows that the speed-curvature power law can be achieved without explicitly optimizing jerk or smoothness either in the planning phase or online.

However, the robot did not accurately follow the path component of the reference. It then seems that the emergence of the power law comes from the *failure* of the robot to accurately follow the high-frequency non-power law position reference, which is interesting. This also seems to be a consequence of low-pass filtering the reference signal by the robot arm system. The position references in x and y dimensions have high-frequency components that get filtered out, leaving single-frequency sinusoid fundamentals, conforming to the aforementioned power law. Similar results were obtained in simulations by Gribble and Ostry (1996) and Schaal and Sternad (2001) where low-pass filtering non-power-law input signals produced power law trajectories.

Related to this result, several studies with human participants have shown that it is difficult to accurately track targets that don’t follow the speed-curvature power law (Viviani, 1988; Viviani, Campadelli, & Mounoud, 1987; Viviani & Mounoud, 1990). However, the subjects did accurately follow the path component of the trajectory and the rhythm of the target. We further discuss possible mechanisms in the section “higher levels”.

### 4.3. A second order simple model accounts for the robot’s behavioral features

Human behavior in tracking pseudorandom targets can be accurately modeled by a first-order model with three constant parameters (see review in Parker et al. 2020). The step-response and frequency response in humans is also modeled by second-order models, bang-bang control, surge control, or the Crossover model (compared in Müller et al, 2017) with various tradeoffs in simplicity and accuracy of modeling.

Here, in turn, we modeled the behavior of the robot itself with a second-order system, a mass on a spring with damping, with three constant parameters (**Figure 5**). Once the parameters were estimated, the model closely reproduced robot position and velocity in visual space in the frequency response task, in the step-reference task, and in tracking elliptic references. The model displayed isochrony and bell-shaped speed profiles in the step-reference reaching task and the power law in the ellipse tracking task (**Figure 5**).

The fact that the model captures all these features of robot behavior with just three parameters is surprising given the multi-level control architecture, the nonlinearities in the lower levels, and differences in motors in each joint. This finding points to an interesting property of hierarchical systems: higher-order loops may appear as linear systems regardless of non-linearities at lower levels. Higher levels provide reference signals to the lower-level systems, so the lower systems are part of the ‘plant’ from the perspective of the higher systems. A certain range of variations and nonlinearities in the plant will be hidden in the behavior of the high-level loop.

If the system for tracking a visual target appears to higher levels of the brain just like the robot arm visual control systems appear to us as experimenters, movement control might be relatively simple for the higher brain structures. As postulated by Viviani and Mounoud (1990), all voluntary movement might be a special case of pursuit tracking, where the only difference from conventional tracking is that the target is internal. The hypothetical higher-level system would only need to specify the virtual target, which is identical to the position reference.

### 4.4. Higher levels of control

In optimal feedback control theory, as well as in industrial robotics control, the solution to the problem of producing a trajectory might involve forward or inverse models, online optimization with a changing horizon, and similar schemes. Those methods are very powerful, especially when coupled with modern computers, precise actuators, and relatively noise-free environments. However, from an academic perspective, they are criticized for not being empirically refutable or biologically plausible (Scott, 2012; Feldman 2015; Powers 2008). In the framework of hierarchical perceptual control (Powers 1973), higher levels should be controlling variables more abstract than lower levels, and also work more slowly having a larger time constant and longer transport delays.

One hypothesis arising from the straight and curved movements of the robot arm is that the present architecture is missing higher levels of control. In the present architecture, the visual position references are set by the experimenter. A hypothetical higher level would be taking the role of the experimenter and would attempt to control or maintain a high-level variable at a desired value by using the position reference as the manipulated variable (much like the current visual position control loops manipulate references to proprioceptive variables to bring visual position to the desired value). The visual position reference would not necessarily be always equal to the produced visual position.

Harris and Wolpert (1998) show that minimizing the variance of the hand trajectory from the desired path over a set of movements (accuracy) for a given speed, or minimizing the speed for a given desired accuracy, reproduced the bell-shaped speed profiles and the 2/3 power law. Their result was achieved in the framework of planning an optimal trajectory and executing it in an open-loop manner; however, we propose that their cost functions for speed and accuracy might be treated as explicit higher-order controlled variables that set visual position references. This would amount to independent control of path accuracy and average speed.

Alternatively, elliptic or straight-line movement can be produced rhythmically. There is some evidence that control of movement amplitude and frequency are developed independently. For instance, 5-year-old children occasionally produce sinusoidal movements that match the amplitude but not the frequency of the target, while other children match the frequency but not the amplitude (Mounoud et aI., 1985). This might suggest a closed-loop pattern generator, similar to a phase-locked loop, that produces a patterned reference for the position control system (see Matic & Gomez-Marin 2019). More research is needed to further elucidate these questions.

### 4.5. Controlled variables in a hierarchy explain adaptivity to perturbations

The ability of humans and animals to achieve the same task outcome using different motor means has been termed as the phenomenon of *motor equivalence*. The problem it poses to motor control theories is the apparent rapid selection of correct-enough means from the space of all possible means. While motor synergies and hierarchical control are proposed as the solution for the problem (Bernstein, 1967), the concept of synergy in motor control literature is defined in many different ways (see review by Bruton and O’Dwyer, 2018). Our reach control system fits the definition of Latash et al (2007), and we termed it the *reach synergy*.

To probe the system, we blocked the wrist of the robot and put it through the same battery of tests as in the normal condition: the step reference reaching task and tracking pseudorandom and elliptic targets. Without any reprogramming or autonomous learning algorithms, the robot still performed the task with similar performance to the normal condition (**Figure 6**), with the exception of pen angle not being controlled (as the pen was fixed perpendicularly to the hand, and the wrist was blocked). Wrist blocking was also modeled in the DIRECT model (Bullock et al., 1993) where they argued that fast adaptations to losing a degree of freedom probably exclude complex planning as a relevant mechanism as it would take too much time, and the same effect can be achieved by simpler schemes. Our result seems to be consistent with the *minimum intervention principle* (Todorov and Jordan, 2002b) where the task-level variable of reaching toward a goal is maintained, and the variability caused by blocking the wrist is taken up by task-irrelevant variables of elbow and shoulder joint angles. The minimum intervention principle may emerge from an online movement optimization algorithm, however, here we achieve the same result without optimization, by having a flexible control hierarchy.

Maintaining pen pressure and angle are indeed important skills in handwriting. Measures of quality of control have been linked, for instance, to dysgraphia as a diagnostic criterion (Mekyska et al 2017). Modeling contact forces in model-based control and optimal feedback control is still an open problem. Control systems for pen pressure and pen angle were implemented for this robot as slow, but precise systems in the higher level. We tested these two control systems by tilting the graphics tablet by 30° and keeping the reference for the pen angle toward the tablet at 0° and the reference for pressure at 50%. Next, the robot performed the battery of tracking tests (**Figure 7**), and we found the performance close to the normal condition. The robot automatically adjusted the height of the tip of the hand and the angle of the wrist in order to maintain the angle and pressure references. In this case, we can see that precise pressure control is crucial for maintaining the pen on the tablet, similarly to human handwriting.

The visual coordinate system was “retina-based” in the sense that the two-dimensional visual field recorded by the camera was the working space of the robot. It was somewhat primitive, as it could only find the location of a green marker in two dimensions. However, it was robust to perturbations. The robot arm and the camera, and their respective proprioceptive and visual coordinate systems were only roughly aligned to begin with, and this was sufficient for normal operation. In the first test, we extended the arm with a 12 cm long piece of plastic and put a marker on the tip of the plastic instead of the tip of the hand, simulating writing with a longer pen or reaching with a stick. The visual system had no information about the size of the stick, or for that matter, the size of the robot arm or the configuration of its angles, but only the location of the marker. This was sufficient to enable the robot to track the reference with this ‘tool’.

In the second test, we rotated the camera by 30°. This made the relationship between visual location proprioceptive location variables more nonlinear than in normal operation. The performance in this test was somewhat worse than in normal condition (**Figure 7**), but the tasks were still successfully performed: in the step-reference condition, the hand tip reached the reference position and settled at that position, and in target tracking, the hand tip followed the reference signal in a similar manner to the normal situation.

These perturbations to the visual system do not greatly affect performance because there are no explicit coordinate transformations between the visual and kinesthetic loops. All transformations are implicit: the higher levels tell the lower levels to ‘move until the higher-level reference state is achieved’. Moving in approximately the right direction seemed to be enough. As discussed, most nonlinearities in the lower levels were absorbed by the high-gain higher-level control systems, at least in the low frequency, low-speed movement.

The geometric and kinematic definitions of controlled variables used in the arm were selected and adapted to fit with this specific robot arm, largely based on the previous computer simulations. While we suspect similar variables might be found in human arm control, we have no direct evidence to support the claim. Following the performance of the robot arm in this study, we suggest that an architecture featuring hierarchical arrangement of controlled variables might be a plausible solution for biological arm control.

### 4.6. Limitations and perspectives

A limitation of the present study is the lack of direct literal comparison of robot behavior to human behavior in the same tasks, and instead comparing invariances and trends. The mechanical and sensory properties of the arm were not on par with the human sensory-motor system to allow such a comparison. With the aim of creating a higher fidelity model of a visually, tactually, and proprioceptively controlled human arm, the improvements would make the arm slightly faster and the sensors more numerous, but maybe not more precise. The improvements would not remove transport delays or noise, because those properties are present in biological arm control systems.

Mechanically, backlash in the geartrain of the motors (also known as slop or play) seems to be a major obstacle for human-like movement, as it puts a hard limit on the precision and bandwidth of the system that cannot be improved by higher quality sensors. With all the nonlinearities, slowness, and fatigability in human muscles, human joints are backlash-free. Therefore, a higher fidelity model should put an emphasis on removing the backlash from the joints, perhaps by tendon-driven actuators.

The visual system of the present robot is a crude approximation of the human visual system’s object detection in two dimensions. Accurate modeling of visual delays should be maintained, but the resolution and refresh rate could be improved, as well as adding stereo vision for three-dimensional localization. Improvements in the same direction could be made to proprioceptive and haptic sensory systems. In sum, such improved systems would allow testing hypotheses of lower, spinal-level sensorimotor loops, and their interaction with higher-level visual or proprioceptive loops, multi-sensory integration etc.

Additionally, as in studies of human movement, the results are influenced by low-pass filtering the data in the analysis stage, and should be taken with some reserve.

### 4.7. Conclusion

This research has shown that in a robot arm system with a hierarchical control architecture based on simulations by Powers (1999, 2008) several features characteristic of biological movement naturally emerge. The robot is robust to noise, delays and some nonlinearities. We found isochrony and bell-shaped velocity profiles in straight reaching movements and the speed-curvature power law in the fast drawing of ellipses. We showed how they can be achieved without trajectory planning, learning or online optimization. We also showed that a hierarchy of controlled variables can produce a motor equivalence phenomenon, where the robot performs the same visual task either with the wrist freely moving or with the wrist blocked. The system also adapts to different angles of the graphics tablet tilt by relying on pressure and pen angle control. Moreover, the system adapts to extending the arm with a tool, and to rotations of the visual field. Overall, we have demonstrated that our 4 DOF robot arm recapitulates important features of human movement and therefore presents an appealing platform upon which to build and test further models of adaptive behavior, while providing insight into feasible mechanisms of human arm control.

## Author contributions

PV and AM conducted a preliminary study. AGM and AM redesigned the project in its current form. AGM supervised the research. AM built the arm system, wrote the control software, did the experiments, analyzed the data, and performed the modeling. AM wrote the first complete manuscript draft. AM and AGM edited the final version of the manuscript. All authors approved the final version of the manuscript.

## Funding

This work was supported by the Spanish Ministry of Science (pre-doctoral contract BES-2016-077608 to AM; Severo Ochoa of Excellence program SEV-2013-0317 start-up funds to AGM; Ramón y Cajal grant RyC-2017-23599 to AGM).

## Conflict of interests

The authors declare that the research was conducted in the absence of any commercial or financial relationships that could be construed as a potential conflict of interest.

## Acknowledgments

We thank Javier Alegre-Cortés, Regina Zaghi-Lara, Roberto Montanari, Max Parker, and especially Kevin Caref for valuable comments on the manuscript. We are grateful to Regina Zaghi-Lara for photos of the robot arm.

## Notes

### Competing Interest Statement

The authors have declared no competing interest.

